# Genomic analyses identify significant biological processes in DDX21-mediated colorectal cancer cells

**DOI:** 10.1101/2022.04.28.489934

**Authors:** Xueying Wang, Donghong Zhang, Mengshan Wang

## Abstract

Colorectal cancer is the third most common cancer in the US. There has been an incline in the number of young patients with colorectal cancer due to unclear reasons at this point in time. Currently, DEAD-box RNA helicase protein DDX21 is identified as a prognosis marker for early-stage colorectal cancer. However, the mechanism of DDX21 mediated-colorectal cancer is still unknown. Here, our objective is to determine the key molecules and signaling by analyzing the RNA-seq data. The GSE184726 was created by the Illumina NovaSeq 6000 (Homo sapiens). The KEGG and GO analyses indicated Neuroactive ligand−receptor interaction and Ras signaling pathway were the key signaling pathways during the knockdown of DDX21 in colorectal cancer. Moreover, we identified several interactive genes including PTPRC, FN1, ITGAM, RAD51, TRAF6, CCNB1, FOXP3, CCNA2, HIST2H2AC, and HSPA5. Our study may provide new insights into the treatment of colorectal cancer.

## Introduction

Colorectal cancer is one of the leading causes of cancer-related death in the world^1^. However, mortality associated with colorectal cancer has decreased in the past decades, which can be accredited to widespread screening programs, improved medical conditions, and effective systematic therapies^2^. After targeted agents like inhibitors of endothelial growth factor and epidermal growth factor receptor were included in the chemotherapies, the survival rate increased to 30 months^3^. Based on these findings, a number of anti-angiogenic agents such as monoclonal antibodies are approved for clinical use^4^. The evolution of targeted therapies in colorectal cancer has been identified by the gradual recognition of many biomarkers, and recent advances in our knowledge that tumors evolve under the pressure from the treatment^5^.

DEAD-box RNA helicase 21 (DDX21) belongs to the DExD/H box family, which owns the ATP-dependent double-stranded RNA^6^. DDX21 is a multi-functional protein that can regulate ribosomal RNA biogenesis, transcription, RNA metabolism, and the immune response during virus infections^7^. Moreover, DDX21 is highly expressed in a variety of cancers such as breast cancer, colorectal cancer, and gastric cancer. However, the mechanism of DDX21 in colorectal cancer is still unclear^8^.

In this study, we compared and analyzed colorectal cancer cells with the knockdown of DDX21 by using the RNA-seq data. We determined several DEGs and significant signaling pathways. We then obtained the gene enrichment and protein-protein interaction (PPI) network analysis to construct the interacting map and key genes. These genes and biological processes may provide new knowledge of colorectal cancer cells for clinical treatment.

## Methods

### Data resources

Gene dataset GSE184726 was downloaded from the GEO database. The data was created by the Illumina NovaSeq 6000 (Homo sapiens) (Nanjing University, Xianlin Street NO.163, Nanjing 210000, China). The analyzed dataset includes two negative controls and two HCT8 cells with the knockdown of DEAD-box RNA helicase 21.

### Data acquisition and processing

The data were conducted by the R package as previously described^9-16^. We used a classical t-test to identify DEGs with *P* < 0.05 and fold change ≥ 1.5 as being statistically significant.

The Kyoto Encyclopedia of Genes and Genomes (KEGG) and Gene Ontology (GO) KEGG and GO analyses were performed by the R package (ClusterProfiler) and Reactome. *P* < 0.05 was considered statistically significant.

### Protein-protein interaction (PPI) networks

The Molecular Complex Detection (MCODE) was used to construct the PPI networks. The significant modules were produced from PPI networks and String networks (https://string-db.org/). The biological processes analyses were performed by using Reactome (https://reactome.org/), and *P* < 0.05 was considered significant.

## Results

### Identification of DEGs in colorectal cancer cells with the knockdown of DDX21

To determine the effects of DDX21 on colorectal cancer cells, we analyzed the RNA-seq data of HCT colorectal cancer cells with the knockdown of DDX21 by using siRNA. A total of 744 genes were identified with a threshold of *P* < 0.05. The top up- and down-regulated genes were shown by the heatmap and volcano plot (Figure 1). The top ten DEGs were selected in Table 1.

**Figure 1.**
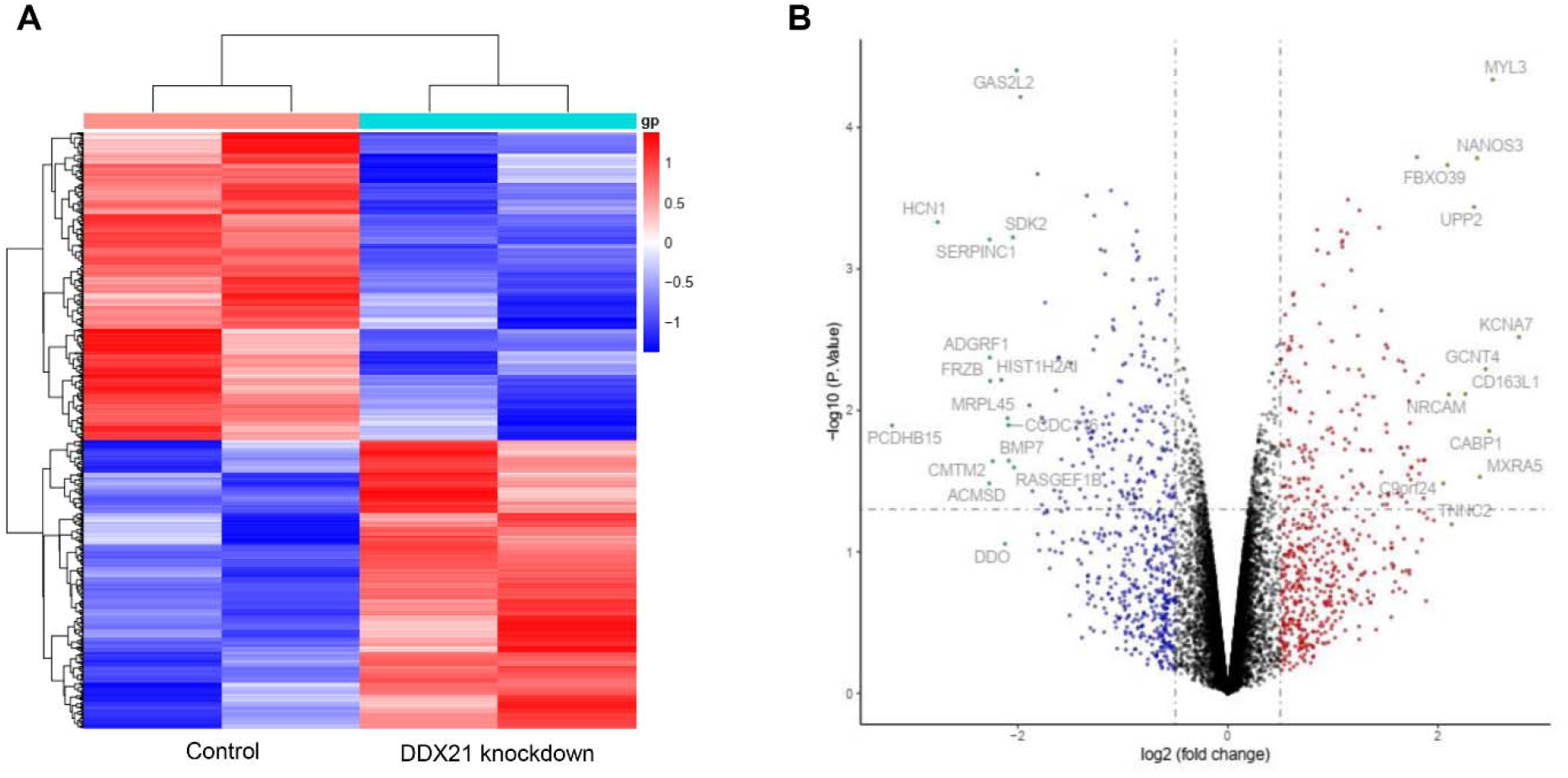
Heatmap and volcano plot in colorectal cancer cells with the knockdown of DDX21. (A) Significant DEGs (*P* < 0.05) were used to construct the heatmap. (B) Volcano plot for DEGs plot in colorectal cancer cells with the knockdown of DDX21. The most significantly changed genes are highlighted by grey dots.

**Table 1.**
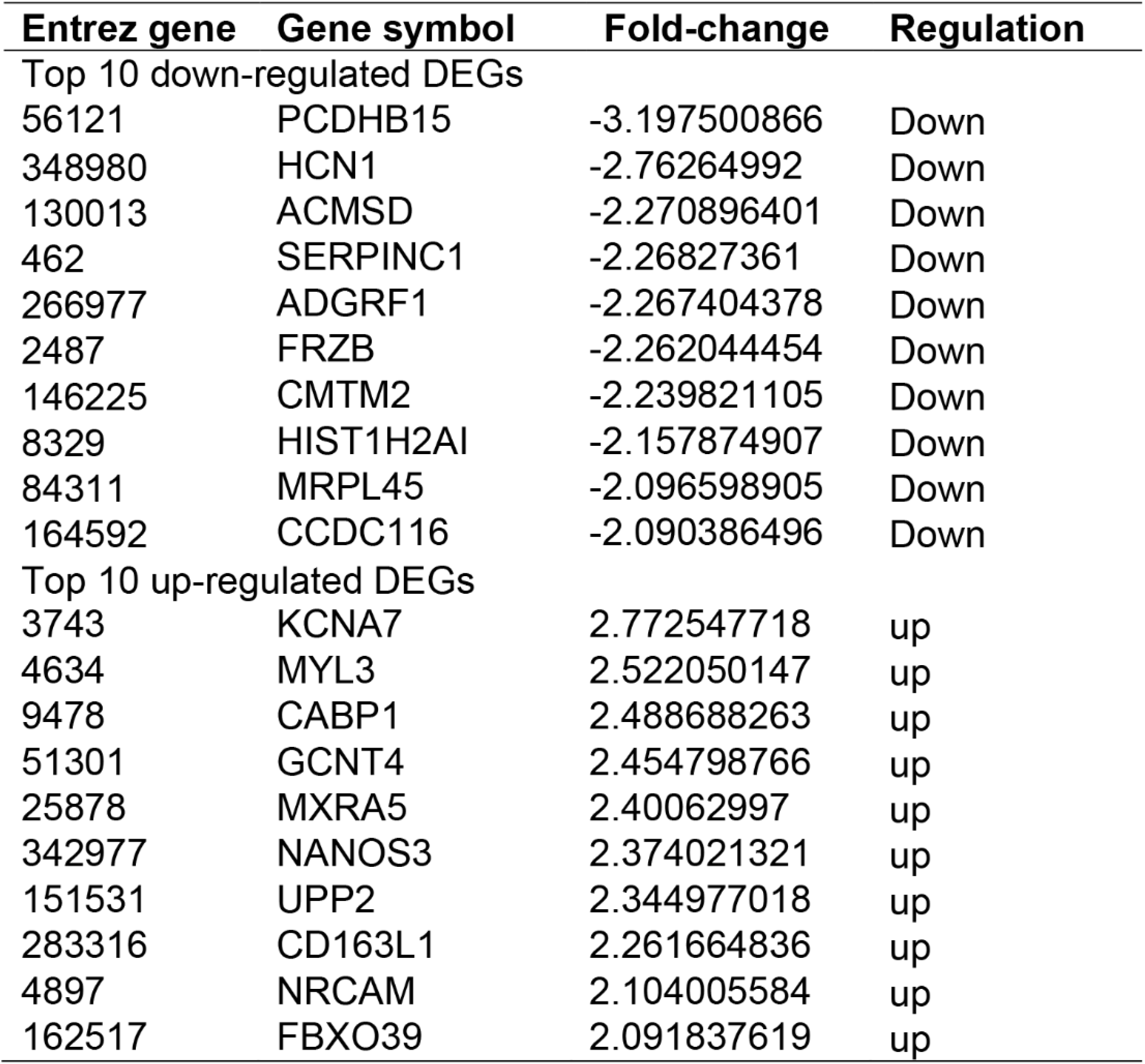

### Enrichment analysis of DEGs in colorectal cancer cells with the knockdown of DDX21

To further study the mechanisms of colorectal cancer cells with the knockdown of DDX21, we performed the KEGG and GO enrichments (Figure 2). We identified the top ten KEGG signaling pathways, including “Neuroactive ligand−receptor interaction”, “Ras signaling pathway”, “Lipid and atherosclerosis”, “NF−kappa B signaling pathway”, “Arachidonic acid metabolism”, “Ether lipid metabolism”, “alpha−Linolenic acid metabolism”, “Linoleic acid metabolism”, “Fat digestion and absorption”, and “Ovarian steroidogenesis”. We identified the top ten biological processes (BP), including “positive regulation of leukocyte migration”, “vitamin metabolic process”, “endocrine process”, “fat−soluble vitamin metabolic process”, “interleukin−2 production”, “response to bacterial lipoprotein”, “regulation of cerebellar granule cell precursor proliferation”, “epithelial cilium movement involved in determination of left/right asymmetry”, “parturition”, and “regulation of toll−like receptor 2 signaling pathway”. We then identified the top ten cellular components (CC), including “apical part of cell”, “apical plasma membrane”, “sarcomere”, “myofibril”, “contractile fiber”, “condensed chromosome”, “I band”, “A band”, “M band”, and “dense core granule”. We identified the top ten molecular functions (MF), including “receptor ligand activity”, “amino acid transmembrane transporter activity”, “amide transmembrane transporter activity”, “proteoglycan binding”, “neutral amino acid transmembrane transporter activity”, “calcium−dependent phospholipase A2 activity”, “ankyrin binding”, “lipopeptide binding”, “polyol transmembrane transporter activity”, and “Toll−like receptor binding”.

**Figure 2.**
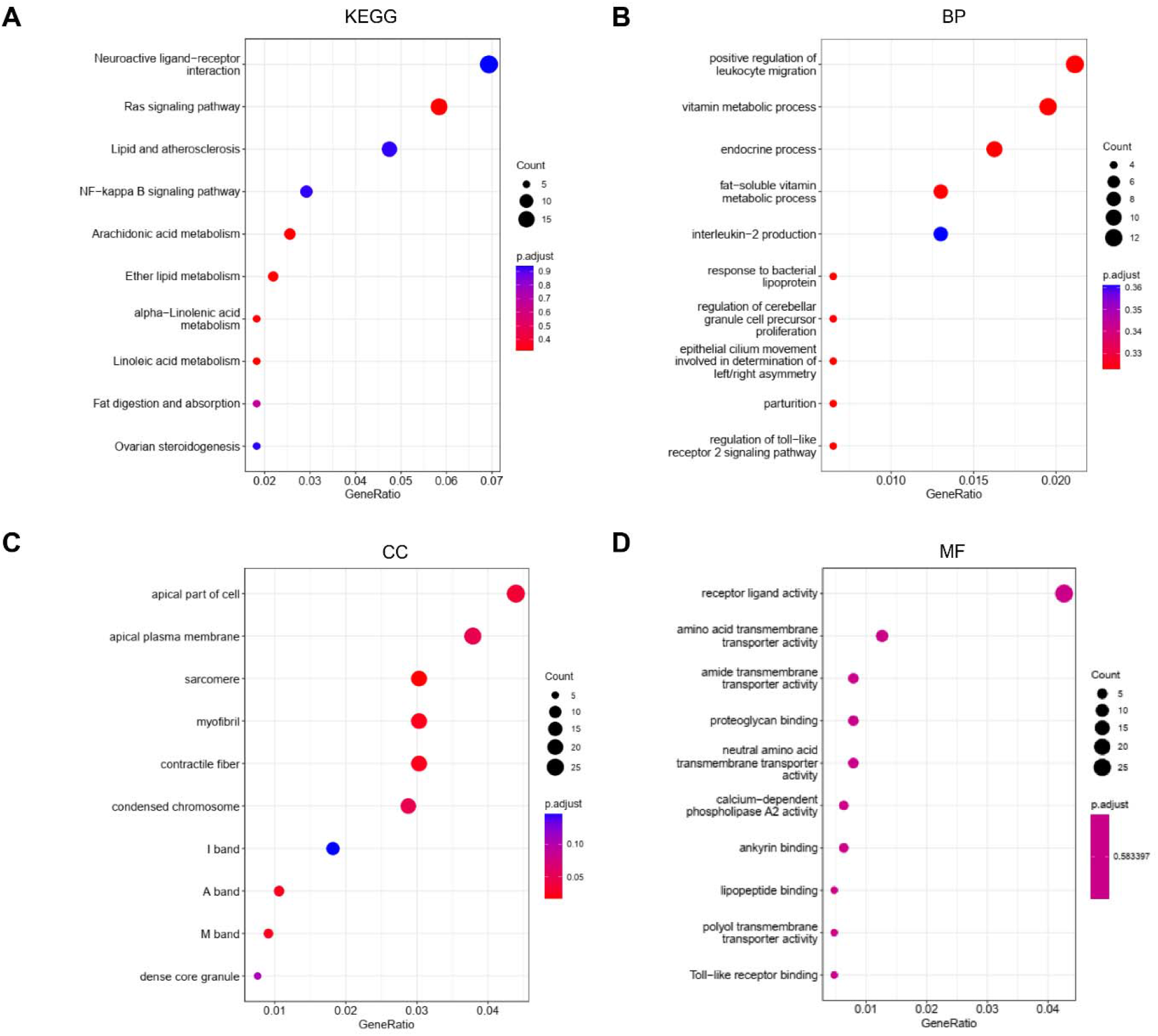
KEGG and GO analyses of DEGs in colorectal cancer cells with the knockdown of DDX21. (A) KEGG analysis (B) BP: Biological processes, (C) CC: Cellular components, (D) MF: Molecular functions.

### PPI network and Reactome analyses

To explore the interactions among the DEGs, we created the PPI network by using 718 nodes and 1,276 edges. The combined score > 0.2 was set as a cutoff by using the Cytoscape software. Table 2 showed the top ten genes with the highest scores. The top two significant modules were selected in Figure 3. We further analyzed the PPI and DEGs with a Reactome map (Figure 4) and identified the top ten biological processes including “Interleukin-37 signaling”, “Regulation of Insulin-like Growth Factor (IGF) transport and uptake by Insulin-like Growth Factor Binding Proteins (IGFBPs)”, “Acyl chain remodeling of PC”, “Acyl chain remodeling of PE”, “RUNX1 and FOXP3 control the development of regulatory T lymphocytes (Tregs)”, “Hydrolysis of LPC”, “Striated Muscle Contraction”, “TP53 Regulates Transcription of Genes Involved in G1 Cell Cycle Arrest”, “Acyl chain remodeling of PS”, and “Downstream signaling of activated FGFR1” (Supplemental Table S1).

**Table 2.** Top ten genes demonstrated by connectivity degree in the PPI network.

| **Gene symbol** | **Gene title** | **Degree** |
| --- | --- | --- |
| PTPRC | protein tyrosine phosphatase receptor type C | 40 |
| FN1 | fibronectin 1 | 39 |
| ITGAM | integrin subunit alpha M | 30 |
| RAD51 | RAD51 recombinase | 29 |
| TRAF6 | TNF receptor associated factor 6 | 28 |
| CCNB1 | cyclin B1 | 27 |
| FOXP3 | forkhead box P3 | 26 |
| CCNA2 | cyclin A2 | 26 |
| HIST2H2AC | H2A clustered histone 20 | 24 |
| HSPA5 | heat shock protein family A (Hsp70) member 5 | 22 |

**Figure 3.**
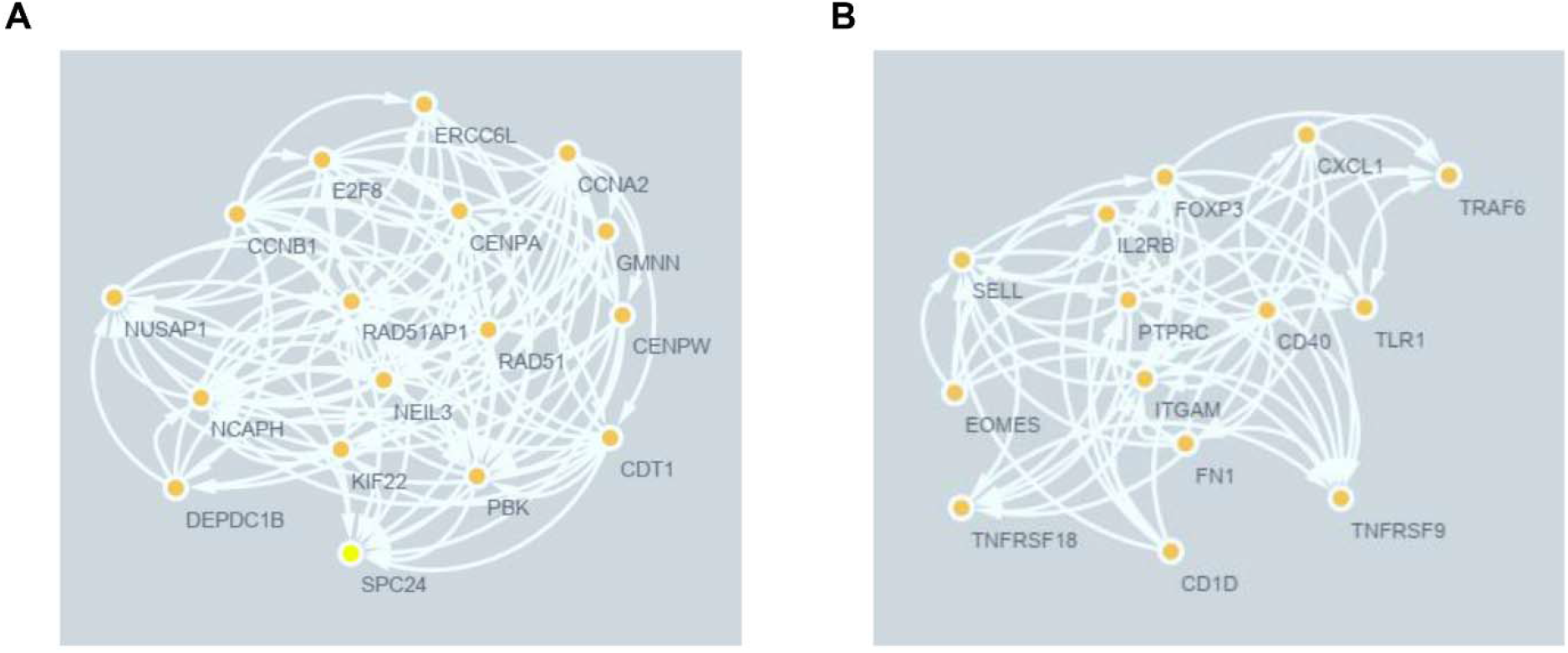
The PPI network analyses of DEGs in colorectal cancer cells with the knockdown of DDX21. The cluster (A) and cluster (B) were constructed by MCODE.

**Figure 4.**
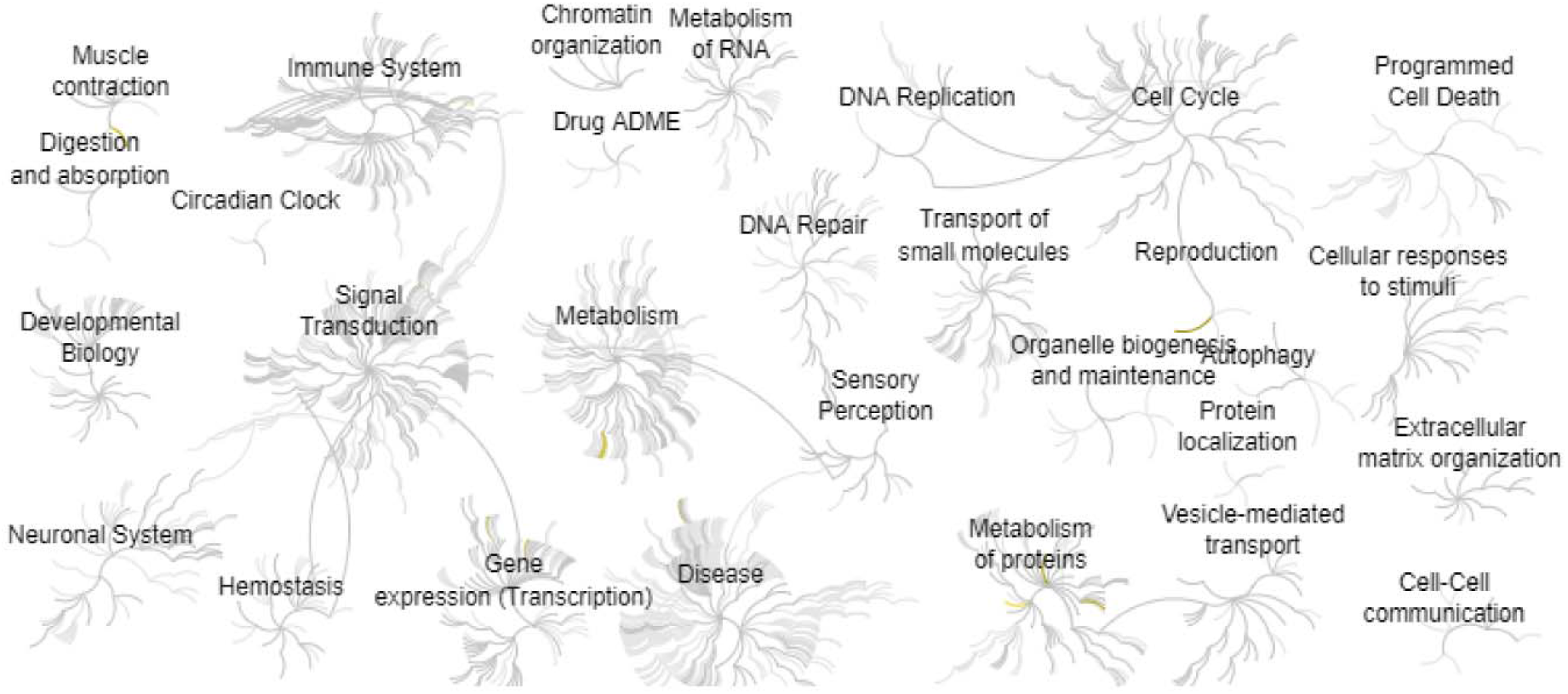
Reactome map representation of the significant biological processes in colorectal cancer cells with the knockdown of DDX21.

## Discussion

The DEAD-box protein family of RNA helicases is important for RNA metabolism, which contains the largest family of RNA helicases^17^. Members of this family share nine conserved motifs that confer RNA binding and RNA unwinding properties^18^. Importantly, they are deregulated in cancers and are critical in the progression of cancers^19^. By analyzing the KEGG and GO enrichments, we found that “Neuroactive ligand−receptor interaction” and “Ras signaling pathway” are the key signaling pathways during the knockdown of DDX21 in colorectal cancer. Similarly, Hui Yao et al found that the Neuroactive ligand-receptor interaction is closely related to the development of colorectal cancer according to the GO and KEGG enrichment analyses^20^. GPCR and RGS signaling pathways are key players in several diseases including cancer, inflammation, and aging-related diseases^21-34^. RAS is a key factor in GPCR signaling. For example, Hiroshi Senoo et al found that Ras GTApases are related to GPCE activation to mTOR signaling^35^. Ian A Prior et al found that Ras is frequently mutated in cancer (10-30%), and Ras may be a key target for treating cancer^36^.

Additionally, we also identified ten key interacting molecules that were involved in the development of colorectal cancer with silent DDX21. Zichuan Liu found that PTPRC acts as a prognostic biomarker for gastric cancer, which is a positive correlation with the progression of gastric cancer^37^. Circadian rhythms refer to oscillations in the various biological processes including inflammation, cancer, aging, and metabolism^11, 38-52^. At the molecular level, such rhythms are created from the transcriptional-translational feedback loops (TTFL) of core clock genes. Interestingly, many of these rhythms are not identical in the tissue-resident PTPRC positive leukocyte population^53^. It is suggested that PTPRC can affect the circadian clock and may regulate the processes of cancer. Lei Dou et al found that the increased CCT3 can enhance cervical cancer progression through FN1^54^. ITGAM was identified as a novel predictor of oral cancer under radiation therapy^55^. Erik Laurini identified that Rad51 is involved in DNA repair and it further regulates the development of cancer^56^. Hua Wu et al found TRAF6 represses colorectal cancer metastasis through CTNNB1 signaling and is targeted for phosphorylation and degradation of GS3B/GSK3b signaling^57^. Er-Bao Chen et al discovered that CCNB1 leads to gastric cancer proliferation and metastasis^58^. Shucai Yang et al found that FOXP2 enhances tumor growth and metastasis by driving Wnt signaling and EMT in lung cancer^59^. Yichao Wang et al found that CCNA2 acts as a potential biomarker of immunotherapy in breast cancer^60^. Fátima Liliana Monteiro found that HIST2H2AC is a new regulator of proliferation and epithelial-mesenchymal transition in breast cancer^61^. Jiewen Fu et al found the expression of HSPA5 is significantly related to the cancer patients with COVID-19^62^.

In summary, this study found the significant DEGs and signaling pathways during the knockdown of DDX21 in colorectal cancer cells. The “Neuroactive ligand−receptor interaction” and “Ras signaling pathway” are the top affected signaling pathways. Our study may provide novel knowledge for the treatment of colorectal cancer.

## Author Contributions

Xueying Wang, Donghong Zhang: Methodology and Writing. Mengshan Wang: Conceptualization, Writing-Reviewing and Editing.

